# Mitotic phosphorylation by Nek6 and Nek7 reduces microtubule affinity of EML4 to alter spindle dynamics and promote chromosome congression

**DOI:** 10.1101/466979

**Authors:** Rozita Adib, Jessica M. Montgomery, Joseph Atherton, Laura O’Regan, Mark W. Richards, Kees R. Straatman, Daniel Roth, Anne Straube, Richard Bayliss, Carolyn A. Moores, Andrew M. Fry

**Affiliations:** Department of Molecular and Cell Biology, University of Leicester, Lancaster Road, Leicester LE1 9HN, U.K.; Institute of Structural and Molecular Biology, Birkbeck College, Malet Street London WC1E 7HX, U.K.; Astbury Centre for Structural Molecular Biology, Faculty of Biological Sciences, University of Leeds, Leeds LS2 9JT, U.K.; Centre for Core Biotechnology Services, University of Leicester, University Road, Leicester, LE1 7RH, UK.; Centre for Mechanochemical Cell Biology, Warwick Medical School, University of Warwick, Coventry CV4 7AL, U.K.

**Keywords:** EML4, Nek6, Nek7, Nek9, mitosis, microtubules

## Abstract

EML4 is a microtubule-associated protein that promotes microtubule stability. We show here that EML4 is distributed as punctate foci along the microtubule lattice in interphase but exhibits reduced association with spindle microtubules in mitosis. Microtubule sedimentation and cryo-electron microscopy and 3D reconstruction reveal that EML4 binds via its basic N-terminal domain to the acidic C-terminal tails of α- and β-tubulin on the microtubule surface. The mitotic kinases Nek6 and Nek7 can phosphorylate EML4 N-terminal domain at S144 and S146 in vitro, and depletion of these kinases leads to increased EML4 binding to microtubules in mitosis. An S144A-S146A double mutant not only binds inappropriately to mitotic microtubules but also interferes with chromosome congression. Meanwhile, constitutive activation of Nek6 or Nek7 reduces EML4 association with interphase microtubules. Together, these data support a model in which Nek6 and Nek7-dependent phosphorylation promotes dissociation of EML4 from microtubules in mitosis thereby altering microtubule dynamics to enable chromosome congression.

## INTRODUCTION

Dynamic instability is an essential property of microtubules that allows them to play diverse roles in intracellular trafficking, organelle positioning, cell migration and cell division (Desai & Mitchison, 1997). Furthermore, the capacity to switch between relatively stable and unstable states is key to the reorganization of the microtubule network that occurs between interphase and mitosis, and which enables assembly of the mitotic spindle upon which chromosomes are segregated. Microtubule dynamics are dependent on the intracellular concentration of tubulin, the rate of GTP hydrolysis by the tubulin heterodimers, and the activity of numerous microtubule-associated proteins (MAPs) that bind the tips and lateral surface of microtubules (Akhmanova & Steinmetz, 2015, Bowne-Anderson, Hibbel et al., 2015, Lawson & Carazo Salas, 2013, van der Vaart, Akhmanova et al., 2009).

One family of MAPs that remains relatively poorly characterized is the EMAP-like, or EML, proteins. EMAP (for echinoderm MAP) was first identified in unfertilized sea urchin eggs as the major non-tubulin component of the mitotic spindle (Suprenant, Dean et al., 1993, Suprenant & Marsh, 1987). However, EMLs are highly conserved and have been described in a variety of organisms, including flies, worms and humans (Bechstedt, Albert et al., 2010, Hueston, Herren et al., 2008, Suprenant, Tuxhorn et al., 2000). While association with microtubules has been shown in different systems, their function in regulating microtubule dynamics remains unclear with studies to date suggesting that different family members might contribute to stabilization and/or destabilization of microtubules (Eichenmuller, Everley et al., 2002, Hamill, Howell et al., 1998, Houtman, Rutteman et al., 2007). Nevertheless, a delay in mitotic chromosome congression observed upon depletion of EML3 or EML4 in human cells confirms their importance for cell division and supports roles for members of the EML family in spindle organization (Chen, Ito et al., 2015, Tegha-Dunghu, Neumann et al., 2008).

Six human EMLs have been described (EML1 to EML6), with EML1 to EML4 sharing a similar organization of an N-terminal domain (NTD) of approximately 175-200 residues encompassing a coiled-coil motif and a C-terminal domain of approximately 650 residues consisting of WD (Trp-Asp) repeats (Fry, O’Regan et al., 2016). Crystallographic studies have revealed that the coiled-coil motif forms a homotrimer, referred to as the trimerization domain (TD), while the WD-repeats assemble into two juxtaposed seven-bladed β-propellers, named the TAPE domain for tandem atypical β-propellers in EMLs (Richards, Law et al., 2014, Richards, O’Regan et al., 2015). A strongly conserved HELP (hydrophobic EMAP-like protein) motif located towards the start of the TAPE domain provides key residues at the interface between the two β-propellers necessary for proper folding. The TAPE domain in isolation does not localize to microtubules although it does bind tightly to soluble tubulin. Interdigitation of the two β-propellers generates a curved sheet with a concave and convex surface; the concave surface shares considerable homology between the different EMLs and point mutations in this region disrupt tubulin binding (Richards et al., 2014). Microtubule binding is rather conferred by the NTD and requires both the TD and a basic region that lies between the TD and start of the TAPE domain. Interestingly, EML5 and EML6 lack the NTD but have three contiguous copies of the TAPE domain. Hence, while EML1 to EML4 can assemble into trimeric complexes that together contain three TAPE domains, EML5 and EML6 contain three copies of the TAPE domain encoded within a single polypeptide. However, as they lack the NTD, it remains unclear whether EML5 and EML6 can bind microtubules.

EML proteins have attracted considerable interest from the cancer community since discovery of translocations involving the genes encoding EML1 and EML4. An EML1-ABL1 fusion protein has been identified in T-cell acute lymphoblastic leukaemia although the frequency appears rare (De Keersmaecker, Graux et al., 2005). In contrast, the EML4-ALK fusion protein is present in a significant proportion (~5%) of lung adenocarcinoma patients, as well as breast and colorectal tumours (Lin, Li et al., 2009, Rikova, Guo et al., 2007, Soda, Choi et al., 2007). Both ABL1 (Abelson 1) and ALK (anaplastic lymphoma kinase) are tyrosine kinases and fusion of the C-terminal catalytic domain of the kinases to the N-terminal region of the EMLs leads to constitutive kinase activation as a consequence of TD-mediated oligomerization (Sabir, Yeoh et al., 2017). Intriguingly, different breakpoints in the EML4 gene lead to distinct EML4-ALK variants that are associated with variable disease progression and therapeutic response in different patients (Bayliss, Choi et al., 2016). EML1 is also implicated in the pathology of an inherited developmental brain disorder where point mutations in the EML1 TAPE domain that potentially destabilize the protein cause neuronal heterotopia in both rodents and humans (Kielar, Tuy et al., 2014).

The microtubule cytoskeleton undergoes dramatic reorganization upon entry into mitosis with a switch from long, relatively stable microtubules to short, unstable microtubules. This switch is largely driven through a change in the complement of MAPs associated with microtubules and significant changes in microtubule nucleation capacity (Heald & Khodjakov, 2015). Observations that both sea urchin EMAP and human EML4 undergo phosphorylation during mitotic progression suggest that regulation of EML proteins may contribute to these changes in the microtubule network (Brisch, Daggett et al., 1996, Pollmann, Parwaresch et al., 2006). EML2, EML3 and EML4 were identified in a large-scale proteomic analysis of human Nek6 binding proteins (Ewing, Chu et al., 2007). Nek6 is a cell cycle-dependent serine/threonine kinase that is activated in mitosis, together with the very closely related Nek7 kinase, downstream of another member of this kinase family, Nek9 (Belham, Roig et al., 2003, Fry, Bayliss et al., 2017, Richards, O’Regan et al., 2009, Roig, Mikhailov et al., 2002). Moreover, Nek6, Nek7 and Nek9 are essential for mitotic spindle assembly and, like EML3 and EML4, for efficient chromosome congression (O’Regan & Fry, 2009, O’Regan, Sampson et al., 2015, Tegha-Dunghu et al., 2008, Yin, Shao et al., 2003).

Here, using antibodies against the endogenous protein, we set out to explore how EML4 might be regulated through the cell cycle. We not only show that the microtubule affinity of EML4 is reduced in mitosis but also that this results from phosphorylation by the Nek6 and Nek7 kinases in the N-terminal microtubule-binding region. We present biochemical and structural data to show that association of EML4 with microtubules occurs through electrostatic interactions between a basic region within EML4 and the acidic tubulin C-terminal tails. Finally, we demonstrate that EML4 is a microtubule stabilizing protein and its displacement from spindle microtubules is essential for chromosome congression in mitosis.

## RESULTS

### Association of EML4 with microtubules is reduced in mitosis

To explore whether the subcellular localization of EML4 is regulated in a cell cycle-dependent manner two different EML4 antibodies were used. The first detects the N-terminal region between residues 150 and 200, while the second detects the C-terminus between residues 951 and 981 (Fig. 1A). Antibody specificity was confirmed by detection of a band at the expected size for EML4 (120 kDa) by Western blot analysis and loss of this band upon depletion of EML4 with two distinct siRNA oligonucleotides (Fig. 1B). Likewise, confocal microscopy with these two antibodies revealed cytoplasmic staining that co-localised with microtubules in human U2OS osteosarcoma cells and that was lost upon depletion of EML4 (Fig. 1C, D). Strikingly, comparison of EML4 localization in interphase and mitotic cells with these antibodies revealed that while the protein associated tightly with the microtubule network in interphase, it was only weakly detected on spindle microtubules in mitosis (Fig. 1E, F). Quantification revealed an approximate two-fold reduction in co-localization of EML4 with microtubules in metaphase compared to interphase (Fig. 1G, H). A similar loss of association with microtubules was observed for recombinant EML4 upon transition from interphase to mitosis in U2OS cells stably expressing YFP-tagged EML4 (Fig. 1I, J). However, YFP-EML4 was detected on spindles in cells expressing high levels of recombinant protein supporting the hypothesis that it is the affinity of EML4 for microtubules that is reduced in mitosis.

**Figure 1.**
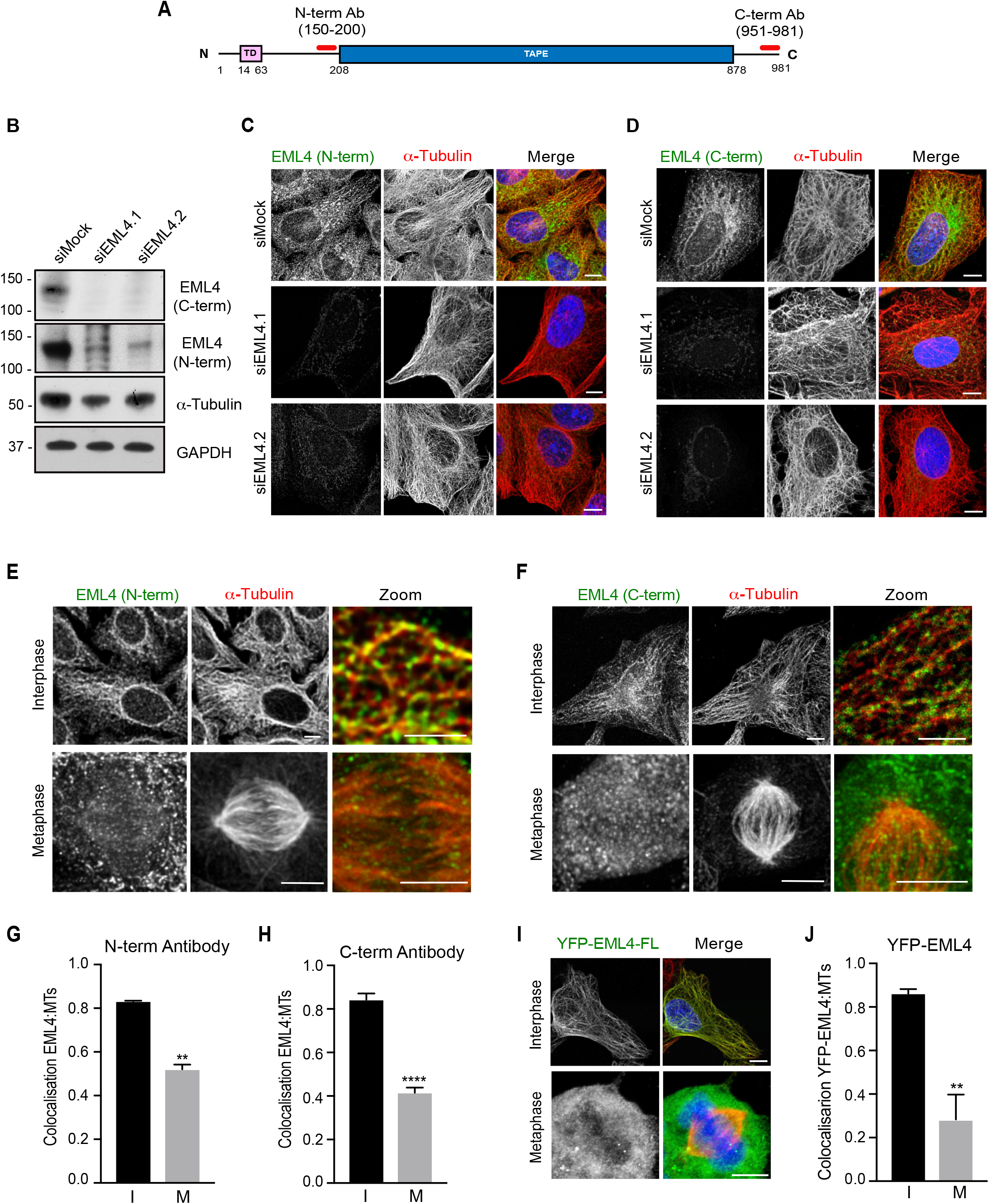
EML4 exhibits reduced affinity for microtubules in mitosis. (**A**) Schematic representation of full-length human EML4 indicating the trimerisation (TD, pink) and TAPE (blue) domains. Epitopes recognised by the two commercial EML4 antibodies (red) map to residue 150-200 (N-term Ab) and 951-981 (C-term Ab). (**B**) Lysates prepared from U2OS cells that were either mock-depleted or depleted of EML4 for 72 h were analysed by Western blot with antibodies indicated. M. wts (kDa) are indicated. (**C & D**) U2OS cells were either mock-depleted or depleted of EML4 for 72 h before being stained with the EML4 antibodies indicated (green) and α-tubulin (red) antibodies. (**E & F**) U2OS cells were stained with the EML4 antibodies indicated (green) and α-tubulin (red) antibodies and imaged by confocal microscopy. Magnified views (zoom) are shown as merges. (**G** & **H**) Co-localization between EML4 and microtubules for cells shown in E & F are shown with the y-axis indicating the mean Pearson’s correlation coefficient (R) from 5 lines per cell in 10 cells (±S.D.). (**I**) U2OS cells were mock transfected or transfected with YFP-EML4-FL 24 h before being stained with GFP and α-tubulin antibodies. DNA was stained with Hoechst 33258 (blue); scale bars, 5 µm. (**J**) Co-localization between the GFP and microtubules for cells shown in G was calculated as in E and F. Scale bars in C, D and G, 5 µm.

### The EML4 NTD is phosphorylated in mitosis

To explore how this cell cycle-dependent change in microtubule affinity is regulated, we first examined whether the endogenous EML4 protein was modified upon mitotic entry. Although Western blot analysis revealed no change in abundance of EML4 between extracts prepared from interphase and mitotic HeLa and U2OS cells, there was a distinct reduction in electrophoretic mobility in mitosis compared to interphase (Fig. 2A). We then tested which region of the EML4 protein was required for microtubule binding and, as previously shown for EML1 (Richards et al., 2015), confirmed that this property is conferred by the NTD (residues 1-207) and not the TAPE domain (residues 208-878) of EML4 (Fig. 2B, C). Interestingly, the isolated EML4 NTD not only co-localised very efficiently with microtubules, but also led to formation of extensive microtubule bundles in the cytoplasm, a phenotype not seen upon expression of the full-length EML4 protein. As microtubule association is dependent on the EML4 NTD, we asked whether a similar cell cycle-dependent change in gel migration was detected upon expression of the isolated NTD. While no change in migration was observed for YFP alone, the YFP-tagged EML4-NTD migrated as two distinct bands in lysates prepared from mitotic cells as compared to a single band in interphase cell lysates (Fig. 2D). This suggests that a substantial proportion (~25-30%) of the EML4 NTD is modified upon entry into mitosis. To confirm that this gel-shift was the result of phosphorylation, we treated lysates taken from mitotic cells expressing the YFP-EML4-NTD protein with λ-protein phosphatase and observed a dose-dependent loss of the slower migrating form confirming that this is a phosphorylated version of the EML4-NTD protein (Fig. 2E). The fact that only a fraction of the recombinant protein is phosphorylated is consistent with ectopic expression of the EML4-NTD saturating the cellular kinase responsible for this modification.

**Figure 2.**
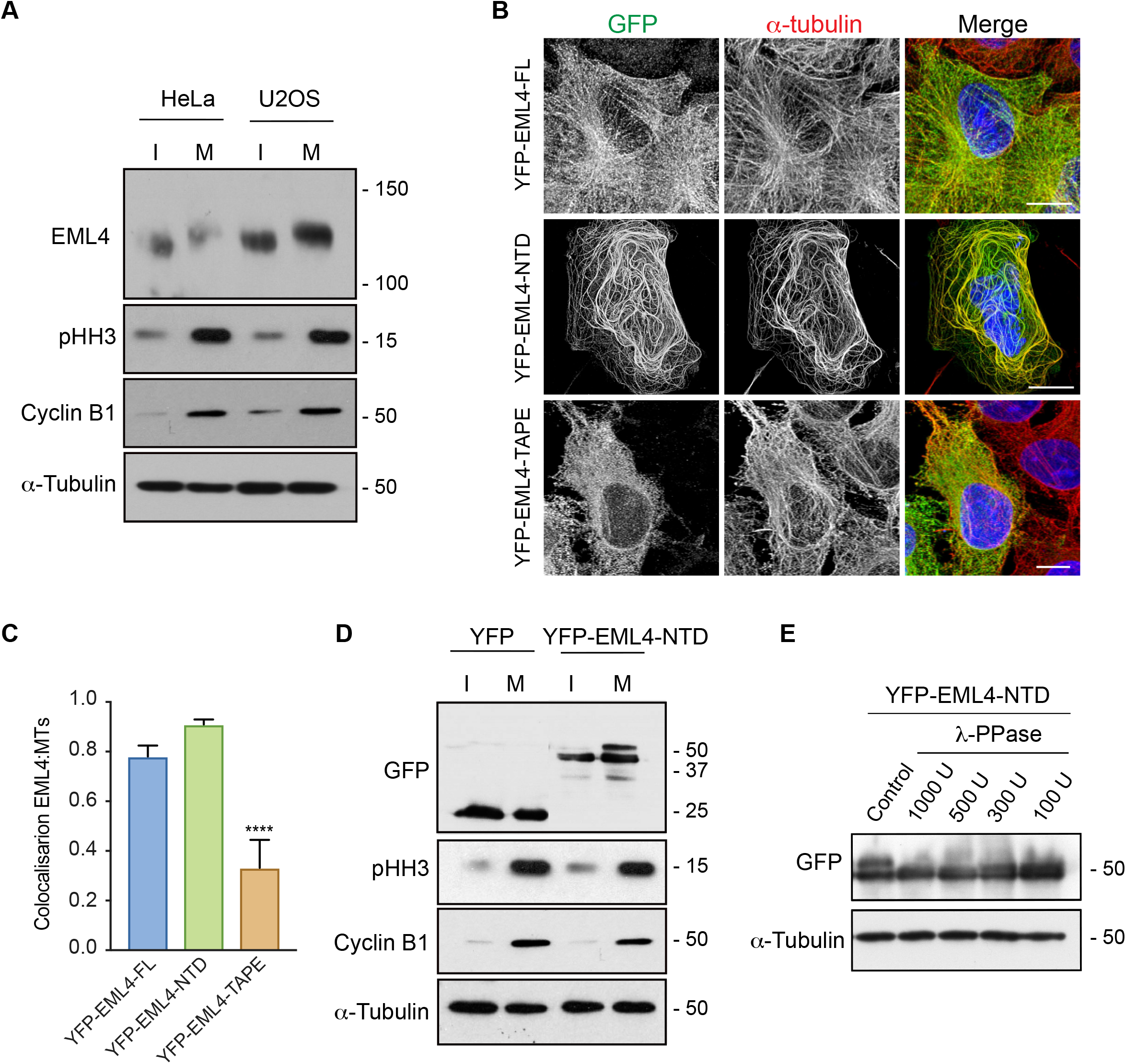
The EML4 NTD binds microtubules and is phosphorylated in mitosis. (**A**) HeLa and U2OS cells were either untreated (I, interphase) or treated with nocodazole for 16 h to arrest them in mitosis (M) before being lysed and analysed via Western blot with antibodies indicated (pHH3, phospho-histone H3). (**B**) U2OS cells were transfected with the EML4 constructs indicated before being fixed and stained with antibodies against GFP (green), α-tubulin (red). DNA was stained with Hoeschst 33258 (blue). Scale bar, 5 µm. (**C**) Co-localization between the GFP and microtubules for cells shown in B was calculated as in Fig. 1. (**D**) U2OS cells were transfected with YFP alone or YFP-EML4-NTD for 24 h before being treated and analysed by Western blot as in C. (**E**) U2OS cells were transfected with YFP-EML4-NTD for 24 h before lysates were treated with λ-PPase (enzyme units indicated) for 30 mins and analysed by Western blot with antibodies indicated. M. wts (kDa) are indicated in A, D & E.

### Phosphorylation of EML4 by Nek6 and Nek7 regulates its microtubule affinity

To determine which kinase(s) may be responsible for this phosphorylation, we took advantage of the previous observation of an interaction between EML4 and the mitotic Nek6 and Nek7 kinases (Ewing et al., 2007). We incubated full-length human EML4 protein purified from insect cells with recombinant Nek6 or Nek7 kinase and ATP. The EML4 proteins were then excised from an SDS-polyacrylamide gel and subjected to mass spectrometry. This revealed four sites that were phosphorylated by Nek6 (S144, T490, T609, S981) and four sites phosphorylated by Nek7 (S134, S146, T609, S981) (Fig. 3A, and Supp. Fig. S1). Analysis of published phosphoproteome data on EML4 via PhosphoSitePlus^®^ revealed clustering of phosphorylation within the NTD and at the extreme C-terminus. Of the three sites within the NTD phosphorylated by Nek6 and Nek7, S134, S144 and S146, the latter two were by far most commonly reported serine/threonine phosphorylation sites based on high throughput proteomic discovery mass spectrometry. Moreover, S144 and S146, but not S134, are conserved across vertebrate species and, in most cases, have a hydrophobic residue at position −3 typical of Nek6 and Nek7 phosphorylation sites (Fig. 3B). We therefore focused on the potential importance of these two sites and generated single S144A and S146A phosphonull mutants, as well as a combined S144/146A double mutant, in constructs expressing YFP-tagged EML4 NTD. Western blot analysis of lysates from transfected U2OS cells revealed no major change in migration of the upper band with the single mutants in cells arrested in mitosis. However, the S144/146A double mutant exhibited a substantially reduced gel-shift confirming that phosphorylation of these two sites is likely to be responsible for the reduced gel mobility in mitotic cells (Fig. 3C). An S144/146A double mutant was therefore also generated in a construct expressing YFP-tagged full-length EML4. Strikingly, while the wild-type EML4 protein did not localise to spindle microtubules, the S144/146A double mutant localised strongly to spindle microtubules (Fig. 3D, E). Hence, we conclude that phosphorylation at S144 and S146 reduces the affinity of EML4 for microtubules in mitosis.

**Figure 3.**
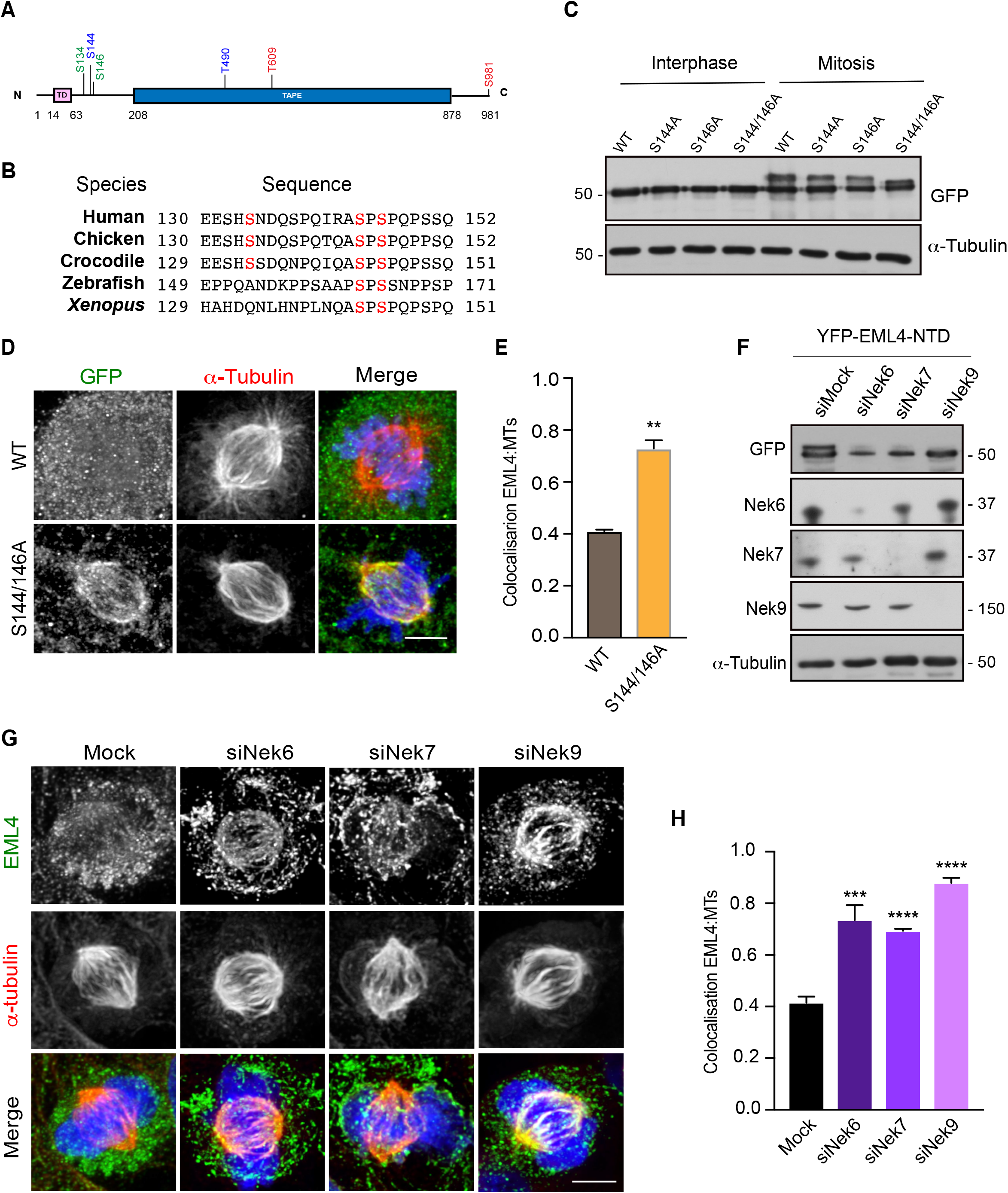
Nek6 and Nek7 regulate association of EML4 with microtubules. (**A**) Schematic representation of full-length EML4 as in Fig. 1A indicating the phosphorylation sites detected by mass spectrometry following incubation in vitro with Nek6 (blue) or Nek7 (green). Sites phosphorylated by both kinases are indicated in red. (**B**) Sequence alignment from species across the five vertebrate classes of the EML4 NTD region spanning the three identified phosphorylation sites (in red). (**C**) U2OS cells were transfected with YFP-EML4-NTD (wild-type, WT, and point mutants) for 24 h before treating with nocodazole for 16 h to arrest cells in mitosis. Cell lysates were analysed by Western blot with antibodies indicated. (**D**) U2OS cells were transfected with YFP-tagged full-length EML4 that was either WT or an S144/146A double mutant for 24 h before being stained with GFP (green) and α-tubulin (red) antibodies. Merge images include DNA stained with Hoechst 33258 (blue). (**E**) The mean Pearson’s correlation coefficient for co-localization between EML4 and microtubules for cells shown in I was calculated from 5 lines per cell in 10 cells ±SD. (**F**) U2OS cells transfected with YFP-EML4-NTD were either mock-depleted or depleted of Nek6, Nek7 or Nek9 48 h before being treated with nocodazole for 16 h to arrest cells in mitosis. Cell lysates were analysed by Western blot with the antibodies indicated. (**G**) Untransfected U2OS cells were either mock-depleted or depleted of Nek6, Nek7 or Nek9 for 72 h before being stained with EML4 (green) and α-tubulin (red) antibodies. DNA was stained with Hoechst 33258 (blue in merge). (**H**) The mean Pearson’s correlation coefficient for co-localization between EML4 and microtubules for cells shown in A was calculated from 5 lines per cell in 10 cells ±SD. M. wts (kDa) are indicated in C & F. Scale bars in D and G, 5 µm;

To determine if Nek6 and Nek7 regulate association of EML4 with spindle microtubules, these kinases were depleted from U2OS cells and the migration of YFP-EML4-NTD analysed in mitotic lysates by Western blot. This revealed that depletion of Nek6 or Nek7, or the upstream Nek9 kinase that is responsible for activating Nek6 and Nek7 in mitosis, led to substantial loss of the slower migrating form of the EML4-NTD protein (Fig. 3F). Furthermore, analysis of endogenous EML4 by immunofluorescence microscopy revealed that depletion of Nek6, Nek7 or Nek9 led to increased association of EML4 with spindle microtubules (Fig. 3G, H). Indeed, depletion of Nek9 led to an increase above that of Nek6 or Nek7 depletion alone. Together, these data suggest that Nek6 and Nek7 act downstream of Nek9 to regulate EML4 localization through phosphorylation of the NTD. To confirm this hypothesis, we expressed constitutively active mutants of Nek6 (Y108A), Nek7 (Y97A) and Nek9 (ΔRCC1) in U2OS cells and found that this led to a significant reduction in association of endogenous EML4 with interphase microtubules (Supp. Fig. S2A, B). These data provide persuasive evidence that the association of EML4 with microtubules is regulated by the Nek9-Nek6-Nek7 kinase module.

### The EML NTD binds microtubules through interaction with tubulin C-terminal tails

In EML1, it was shown that the TD and a basic sequence that lies between the TD and TAPE domain confers microtubule binding to EMLs (Richards et al., 2015). The region between the TD and TAPE domain is also highly basic in the EML4 protein (residues 64-207; pI=10.23) and so we hypothesized that association may depend upon electrostatic interaction between this basic region and the acidic surface of the microtubule created by the C-terminal tails of α- and β-tubulin. These C-termini protrude on the exterior surface of the microtubule and are rich in glutamate (E) residues. A number of MAPs associate, at least in part, via interaction of basic domains with these so-called tubulin ‘E-hooks’, including MAP2 and tau(Manka & Moores, 2018). To seek evidence that the basic nature of the EML4 NTD mediates microtubule association, the tubulin C-terminal tails of microtubules were removed by limited proteolysis with subtilisin. SDS-PAGE analysis confirmed that tubulin now migrated as a doublet following subtilisin treatment indicative of partial cleavage of the C-terminal tails. When incubated with two different concentrations of purified EML4-NTD protein, there was a significant reduction in sedimentation of this protein with the subtilisin-treated microtubules as compared to untreated microtubules (Fig. 4A-C). We planned to use TIRF microscopy of fluorescently labelled proteins to confirm this dependence on the tubulin C-terminal tails. However, it proved impossible to generate a purified version of fluorescently tagged EML4-NTD protein. We therefore undertook this analysis with a purified YFP-EML1-NTD fragment (residues 1-174). Using a mixed population of untreated microtubules labelled with a 561 nm fluorophore and subtilisin-treated microtubules labelled with a 640 nm fluorophore, we found that the EML1-NTD protein bound strongly to untreated microtubules but not to microtubules treated with subtilisin (Fig. 4D, E). Together, these data are consistent with the basic N-terminal region of EML proteins associating with the acidic tubulin C-terminal tails on the microtubule surface.

**Fig. 4.**
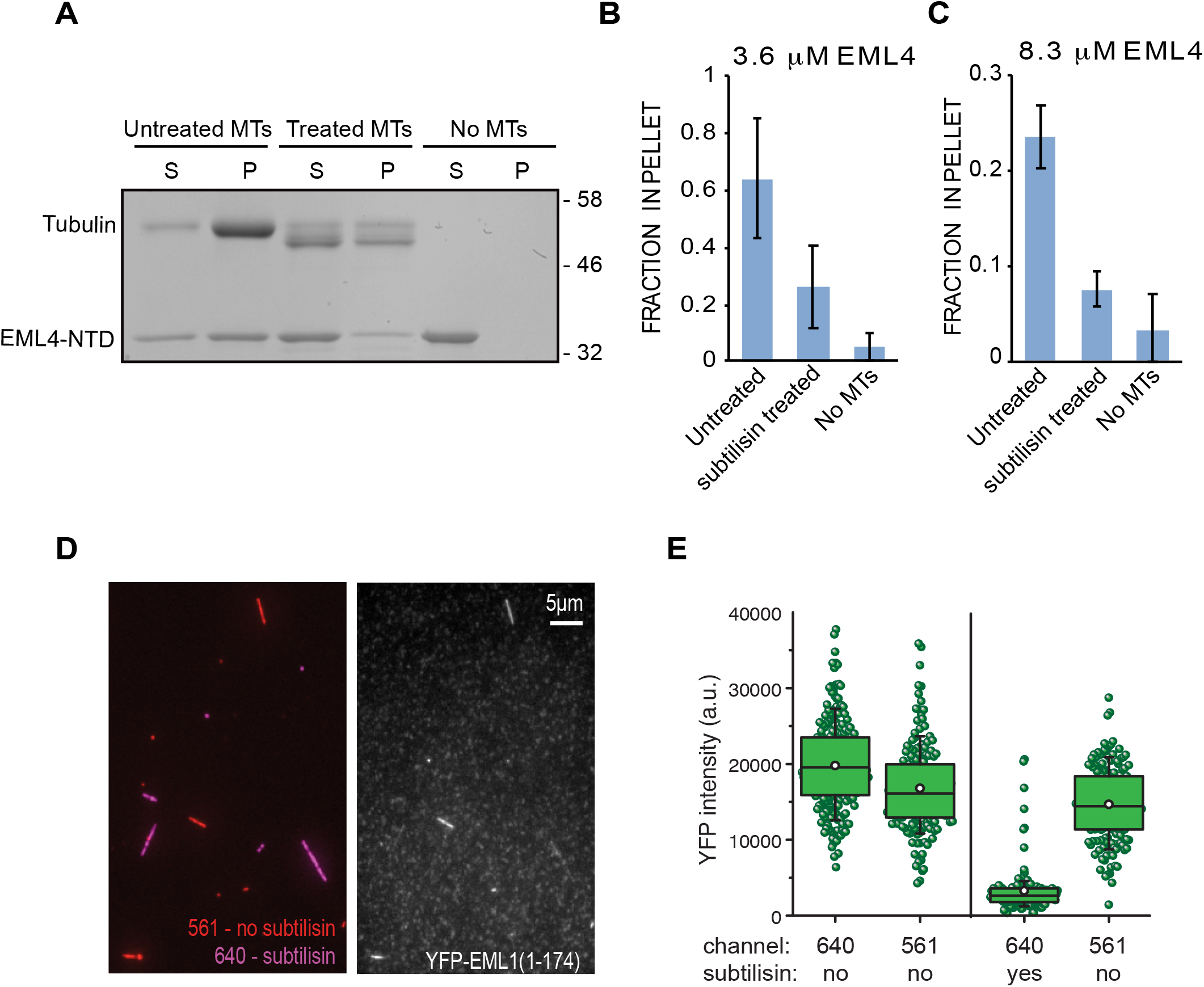
Subtilisin treatment of microtubules leads to loss of association of EML4 and EML1 in vitro. (**A**) Microtubules purified from HeLa cells were either untreated or incubated with subtilisin before re-purification to remove the enzyme. These were then incubated with purified EML4-NTD before sedimentation to generate a supernatant (S) and pellet (P) fraction. Samples were then analysed by SDS-PAGE and Coomassie Blue stain. M. wts (kDa) are indicated. (**B & C**) The mean relative fraction (±SD, n=3) of the EML4-NTD protein present in the pellet fraction from A is shown for two different concentrations of the EML4-NTD protein. (**D**) TIRF image of fluorescently labelled microtubules (left panel) treated with (purple) or without (red) subtilisin before incubation with YFP-EML1-NTD (residues 1-174) (right panel). (**E**) Box plots reveal YFP intensity (a.u., arbitrary units) associated with untreated (561 nm) or subtilisin-treated (640 nm) microtubules. Boxes represent quartiles; whiskers show 10/90% data *±*SD (n>200 microtubules).

### Cryo-electron microscopy reveals binding of EML4 to C-terminal tubulin tails

To examine this interaction of the EML4 NTD with microtubules in more detail, we first performed structured illumination microscopy (SIM) of endogenous EML4 in U2OS cells. This revealed small, evenly sized puncta of EML4 distributed along the cytoplasmic microtubules (Fig. 5A, B). There was no obvious concentration at microtubule ends as has been seen for plus-tip tracking proteins, such as the EBs or ch-TOG proteins (Akhmanova & Steinmetz, 2015). We then used cryo-electron tomography to directly visualize purified EML4-NTD bound to microtubules in vitro. Low-resolution reconstructions showed density corresponding to EML4-NTD bound along the microtubule outer surface consistent with the SIM analysis of endogenous EML4 in cells (Fig. 5C). It also revealed that while there is occasional evidence of evenly spaced interactions with the microtubules every 4 nm (corresponding to tubulin monomers), binding of EML4-NTD is overall rather disordered. We processed images of EML4-NTD bound to MTs using single particle averaging algorithms, and revealed additional density on both α- and β-tubulin due to bound EML4-NTD (Fig. 5D, E, and Supp. Fig. S3). Although the resolution of tubulin in our reconstruction is better than 4 Å (Table 1) and shows clear evidence of discrimination between α- and β-tubulin, density corresponding to EML4-NTD is present at substantially lower resolution. This EML4-NTD density lies above the C-terminal helices H11 and H12 of each tubulin monomer, next to where the C-terminal tails of each monomer emerge from the microtubule wall. This confirms the biochemical evidence for involvement of tubulin C-terminal tails in EML4-NTD binding. The low resolution of the EML4-NTD density and the limited extent of density which is visualized is likely due to the flexible nature of its interaction with the tubulin C-terminal tails, which themselves are unstructured and rarely visualised in cryo-EM reconstructions.

**Figure 5.**
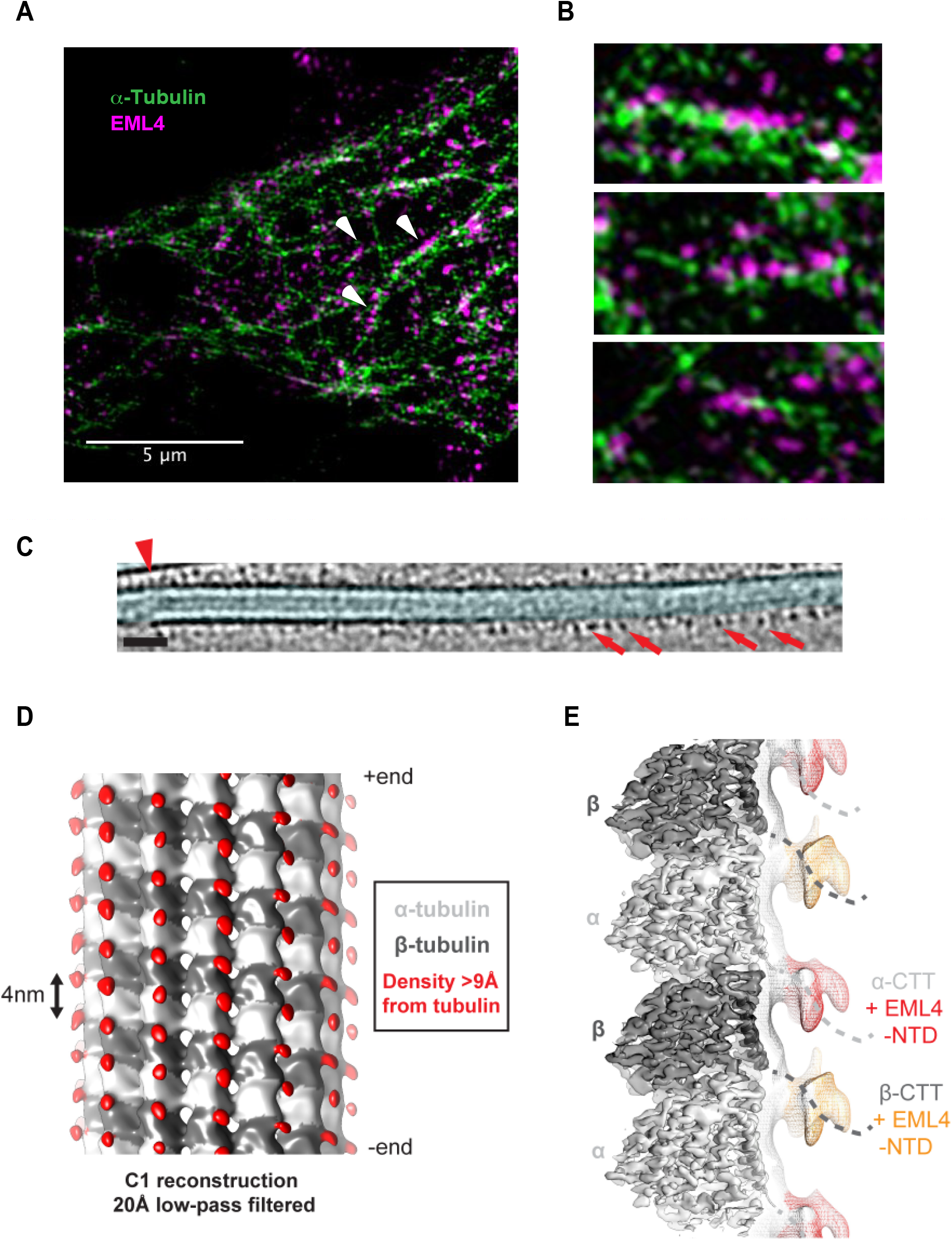
Super-resolution and cryo-electron microscopy reveal EML4-NTD binding microtubules through a flexible interaction with α- and β-tubulin C-terminal tails. (**A**) U2OS cells were stained with EML4 (C-term Ab; magenta) and α-tubulin (green) antibodies and imaged by SIM. (**B**) Magnified views of endogenous EML4 bound to microtubules as taken from A. (**C**) Section through a cryo-tomogram of an EML4 decorated microtubule. The microtubule lumen is indicated in light blue false colour. Red arrows indicate clear microtubule-bound densities, with sizes consistent with EML4-NTD trimers. Red arrowhead: in some regions, a periodicity of 4 nm for extra densities can be observed, consistent with binding to both α- and β-tubulin. Scale bar 30 nm. (**D**) 4.4 Å resolution C1 single-particle cryo-electron microscopy reconstruction of EML4-NTD decorated 13 protofilament microtubules, low pass filtered to 20 Å. Density within 9 Å of the fitted α- and βtubulin atomic models is coloured in light and dark grey respectively, whilst defined, connected density >9Å away is coloured in red. (**E**) 3.6 Å resolution symmetrised reconstruction, showing two tubulin dimers within a single protofilament. Local resolution filtered α-tubulin and β-tubulin density is shown as transparent light and dark grey density respectively. The fitted H12 atomic model is shown as ribbons, with the flexible C-terminal tails indicated by dashed lines. The reconstruction low-pass filtered to 20 Å is also shown as mesh, with extra densities associated with α- and β-tubulin coloured in red and orange respectively.

**Table 1.**
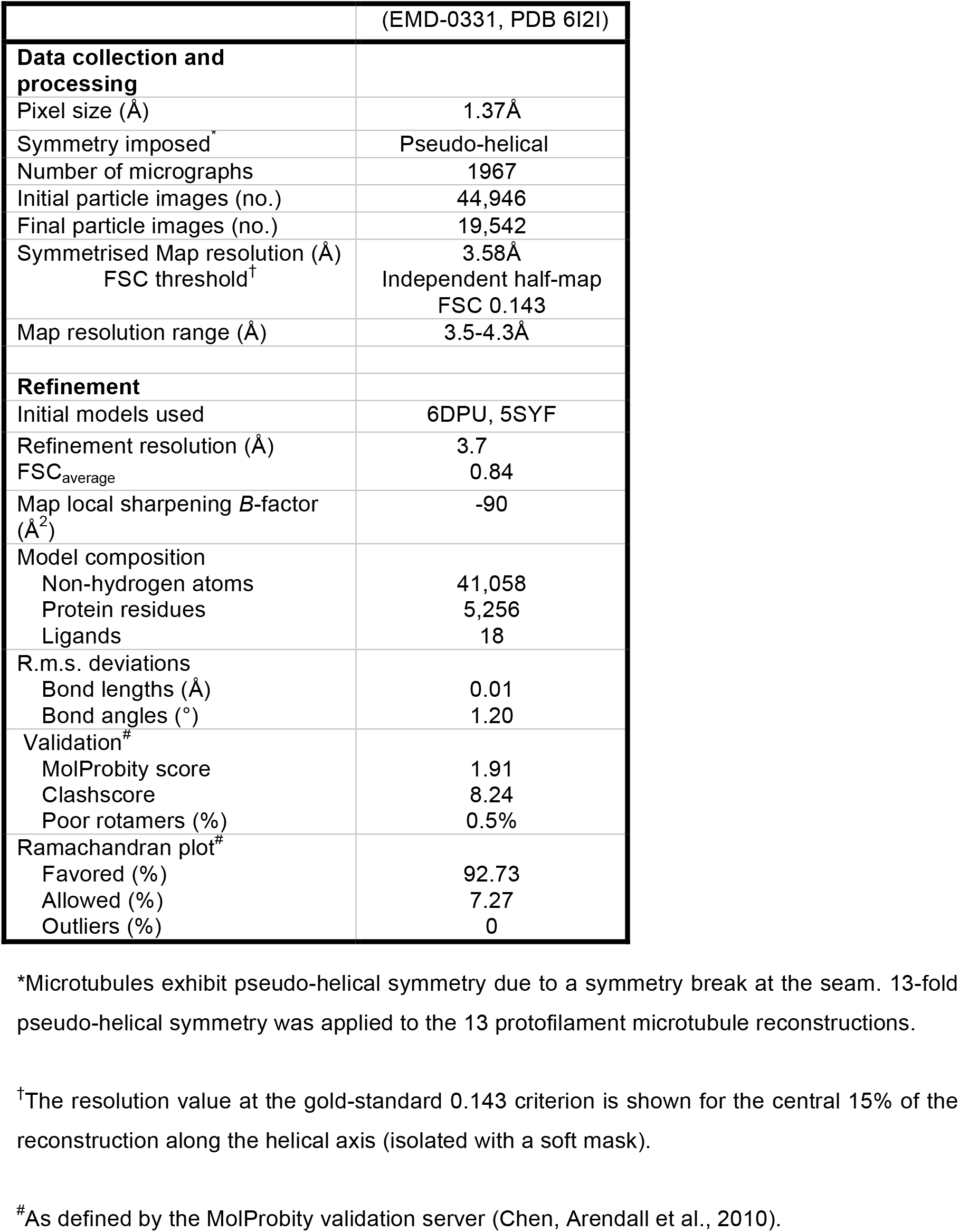
Cryo-EM data collection, refinement and validation statistics.

|  |  |
| --- | --- |
|  | (EMD-0331, PDB 6I2I) |
| **Data collection and processing** |  |
| Pixel size (Å) | 1.37Å |
| Symmetry imposed<sup>*</sup> | Pseudo-helical |
| Number of micrographs | 1967 |
| Initial particle images (no.) | 44,946 |
| Final particle images (no.) | 19,542 |
| Symmetrised Map resolution (Å) | 3.58Å |
| FSC threshold<sup>†</sup> | Independent half-map FSC 0.143 |
| Map resolution range (Å) | 3.5-4.3Å |
| **Refinement** |  |
| Initial models used | 6DPU, 5SYF |
| Refinement resolution (Å) | 3.7 |
| FSC<sub>average</sub> | 0.84 |
| Map local sharpening B-factor (Å<sup>2</sup>) | -90 |
| Model composition |  |
| Non-hydrogen atoms | 41,058 |
| Protein residues | 5,256 |
| Ligands | 18 |
| R.m.s. deviations |  |
| Bond lengths (Å) | 0.01 |
| Bond angles (°) | 1.20 |
| Validation<sup>#</sup> |  |
| MolProbity score | 1.91 |
| Clashscore | 8.24 |
| Poor rotamers (%) | 0.5% |
| Ramachandran plot<sup>#</sup> |  |
| Favored (%) | 92.73 |
| Allowed (%) | 7.27 |
| Outliers (%) | 0 |

<sup>\*</sup>Microtubules exhibit pseudo-helical symmetry due to a symmetry break at the seam. 13-fold pseudo-helical symmetry was applied to the 13 protofilament microtubule reconstructions.
<sup>†</sup>The resolution value at the gold-standard 0.143 criterion is shown for the central 15% of the reconstruction along the helical axis (isolated with a soft mask).
<sup>#</sup>As defined by the MolProbity validation server (Chen, Arendall et al., 2010).

### Displacement of EML4 from microtubules is required for chromosome congression

As microtubules exhibit reduced stability in mitosis, the loss of EML4 from microtubules at this time could be important for these altered dynamics. This would be consistent with a previous report that overexpression of EML4 stabilises microtubule in Cos7 monkey cells (Houtman et al., 2007). To test whether endogenous EML4 promotes microtubule stability in human cells, we first examined the consequences of depleting EML4 in U2OS cells on the sensitivity of microtubules to the depolymerizing agent, nocodazole. Incubation of mock-depleted cells with 75 nM nocodazole did not affect microtubule organization; however, this dose led to loss of an intact microtubule network in EML4-depleted cells (Fig. 6A). Furthermore, EML4 depletion led to reduced sedimentation of microtubules in the presence of 10 µM taxol, also suggestive of impaired stability (Fig. 6B, C). Stable microtubules are subject to more post-translational modifications, including acetylation and detyrosination. However, depletion of EML4 led to significant reduction of both these markers as measured by Western blot or immunofluorescence microscopy (Fig. 6D-F, and Supp. Fig. S4). Together, these data confirm a role for endogenous EML4 in microtubule stabilization and provide a rationale for why it is removed from microtubules upon mitotic entry.

**Figure 6.**
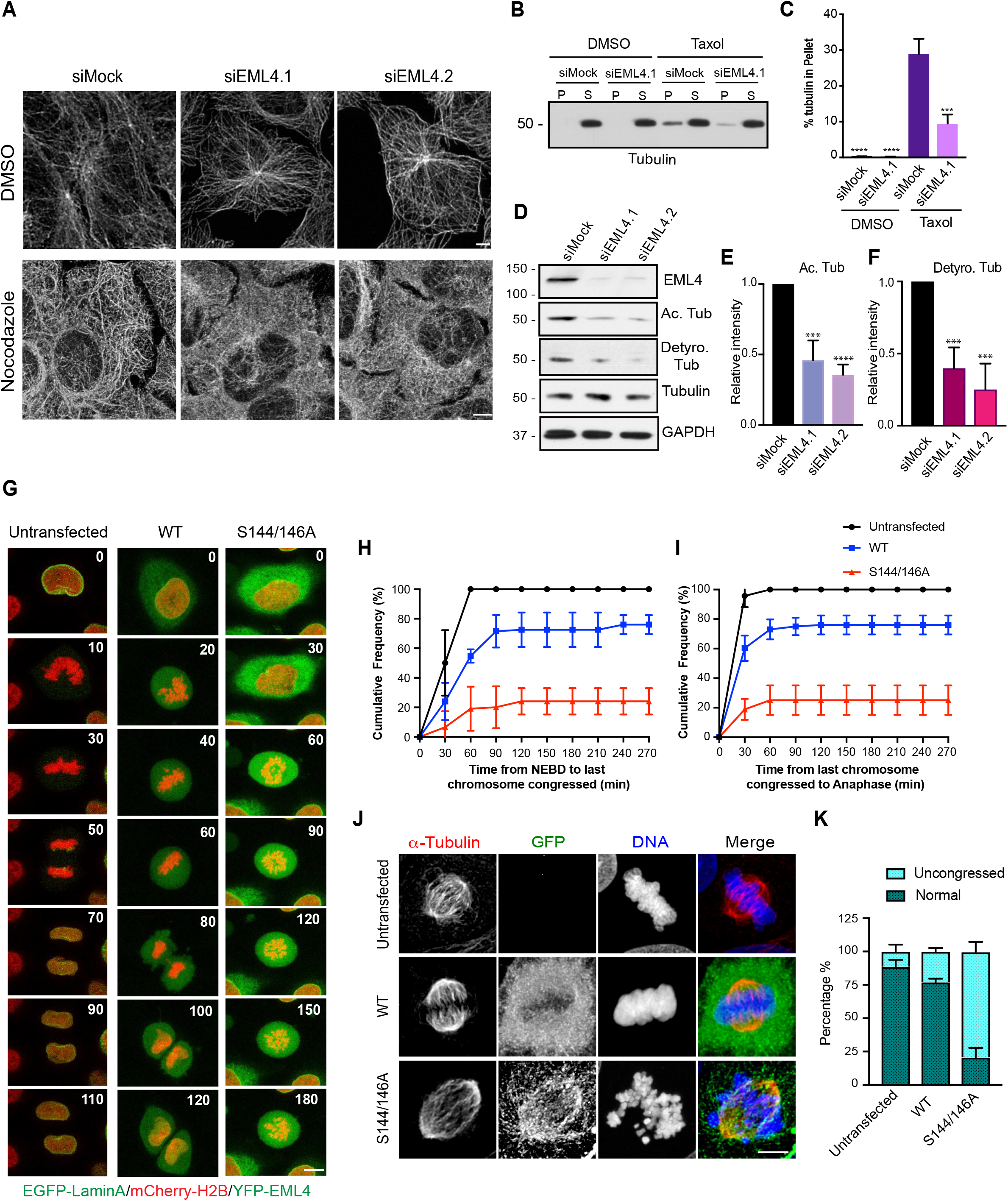
EML4 phosphorylation is required for chromosome congression. (**A**) U2OS cells were either mock-depleted or depleted of EML4 with two different siRNAs for 72 h before being left untreated or treated with 75 nM nocodazole for 2 h. Cells were stained with α-tubulin antibodies. (**B**) Microtubule sedimentation assay performed with lysates prepared from U2OS cells that were either mock- or EML4-depleted before Western blotting the pellet (P) and supernatant (S) fractions for α-tubulin. (**C**) Histogram shows % tubulin in pellet fraction. (**D**) Western blot analysis with antibodies indicated of lysates from U2OS cells that were either mock- or EML4-depleted. (**E & F**) Histogram represents the intensity of acetylated tubulin following EML4 depletion relative to mock-depletion from blots shown in K. (**G**) HeLa:EGFP-LaminA/mCherry-H2B cells were either untransfected (left column) or transfected with YFP-EML4 wild-type protein (WT, middle column) or YFP-EMl4-S144/146A (right column) for 24 hours before time-lapse confocal imaging. Stills from representative movies are shown with time (mins) indicated from mitotic entry; scale bar, 7.5 µm. (**H, I**) Quantification of cells from A indicating time from nuclear envelope breakdown (NEBD) to last chromosome congressed (left graph) and time from last chromosome congressed to anaphase onset (right graph). Data represent means of cumulative frequencies ±S.D.; n=20. (**J**) U2OS cells were either untransfected or transfected with YFP-EML4-FL, WT or S144/146A, for 24 h before being treated with RO-3306 for 16 h, followed by 4 h MG-132 treatment. Then cells were fixed and stained with GFP and α-tubulin antibodies. Merge images include DNA stained with Hoechst 33258 (blue). (**K**) The histogram shows the percentage of cells with normal or uncongressed chromosome in mitosis from at least 30 cells. Scale bars in A, G & J, 5 µm; M. wts (kDa) are indicated in B & D.

To directly test the importance of EML4 displacement from microtubules by phosphorylation in mitosis, we undertook time-lapse confocal imaging of HeLa cells expressing wild-type and a phosphonull EML4 mutant. These cells also stably express an EGFP-lamin A construct that labels the nuclear envelope and an mCherry-histone H2B construct that labels the chromatin allowing visualization of nuclear envelope breakdown and chromosome dynamics. Whereas in untransfected cells, nuclear envelope breakdown (NEBD) was quickly followed by chromosome congression and anaphase onset (as indicated by sister chromatid separation), these events were delayed in cells expressing the wild-type EML4 protein (Fig. 6G-I). Again this can be explained by overexpression of EML4 saturating the machinery required to regulate its localization. Strikingly though, expression of the phosphonull S144/146A double mutant protein led to dramatic failure of both chromosome congression and anaphase onset with less than 30% cells successfully completing cell division after 4 hours in mitosis (Fig. 6GI). Fixed imaging of U2OS cells arrested in mitosis with MG132 to prevent anaphase onset revealed that 90% of untransfected cells and 75% of cells transfected with wild-type EML4 had condensed chromosomes that were fully aligned on the metaphase plate (Fig. 6J, K). In contrast, only 20% of cells transfected with the EML4-S144/146A mutant had fully congressed chromosomes. Moreover, staining for the microtubule network revealed that the spindle microtubules in these cells were unusually long (Fig. 6J). Thus, the kinase-dependent displacement of EML4 from microtubules in mitosis is necessary for assembly of a functional mitotic spindle capable of efficient chromosome congression and cell division.

## DISCUSSION

Human EML4 is phosphorylated on serine/threonine residues in mitosis(Pollmann et al., 2006). Here, we show that EML4 undergoes phosphorylation at S144 and S146 in the N-terminal microtubule-binding region in mitosis and that this reduces its affinity for microtubules. This phosphorylation is catalysed by the Nek6 and Nek7 kinases and perturbs electrostatic binding of the EML4 protein with the tubulin C-terminal tails that extend from the surface of the microtubule lattice (Fig. 7A). Confocal and super-resolution microscopy indicate the presence of EML4 foci that are distributed along the length of polymerised microtubules. However, the molecular details of how EML proteins bind microtubules had remained unclear. Here, we show that this interaction most likely occurs through electrostatic interaction of the basic N-terminal domain of EML4 with the acidic tubulin C-terminal tails (so-called E-hooks) that are exposed on the surface of the microtubule. This conclusion is based not simply on the fact that phosphorylation, i.e. the addition of negative charge, weakens the interaction, but on the observation that limited proteolysis of the microtubules with subtilisin that cleaves off these C-terminal tails, abrogated binding. Indeed, cryo-electron microscopy and image reconstruction provided definitive evidence that the EML4 NTD binds at the site on both α- and β-tubulin where the C-terminal tails emerge. Furthermore, spacing of the EML4-NTD foci suggests that the protein can bind to both tubulin monomers within an individual heterodimer. It will be intriguing to explore whether the acidic tubulin C-terminal tails wrap around a trimeric form of the EML4-NTD or whether the basic region that extends from the trimerization domain forms a hollow structure within which the tubulin C-terminal tail is threaded (Fig. 7B). What is also not yet clear is whether the full-length EML4 protein binds in an identical manner and how the TAPE domain, which binds soluble tubulin heterodimers, is oriented with respect to the microtubule.

**Figure 7.**
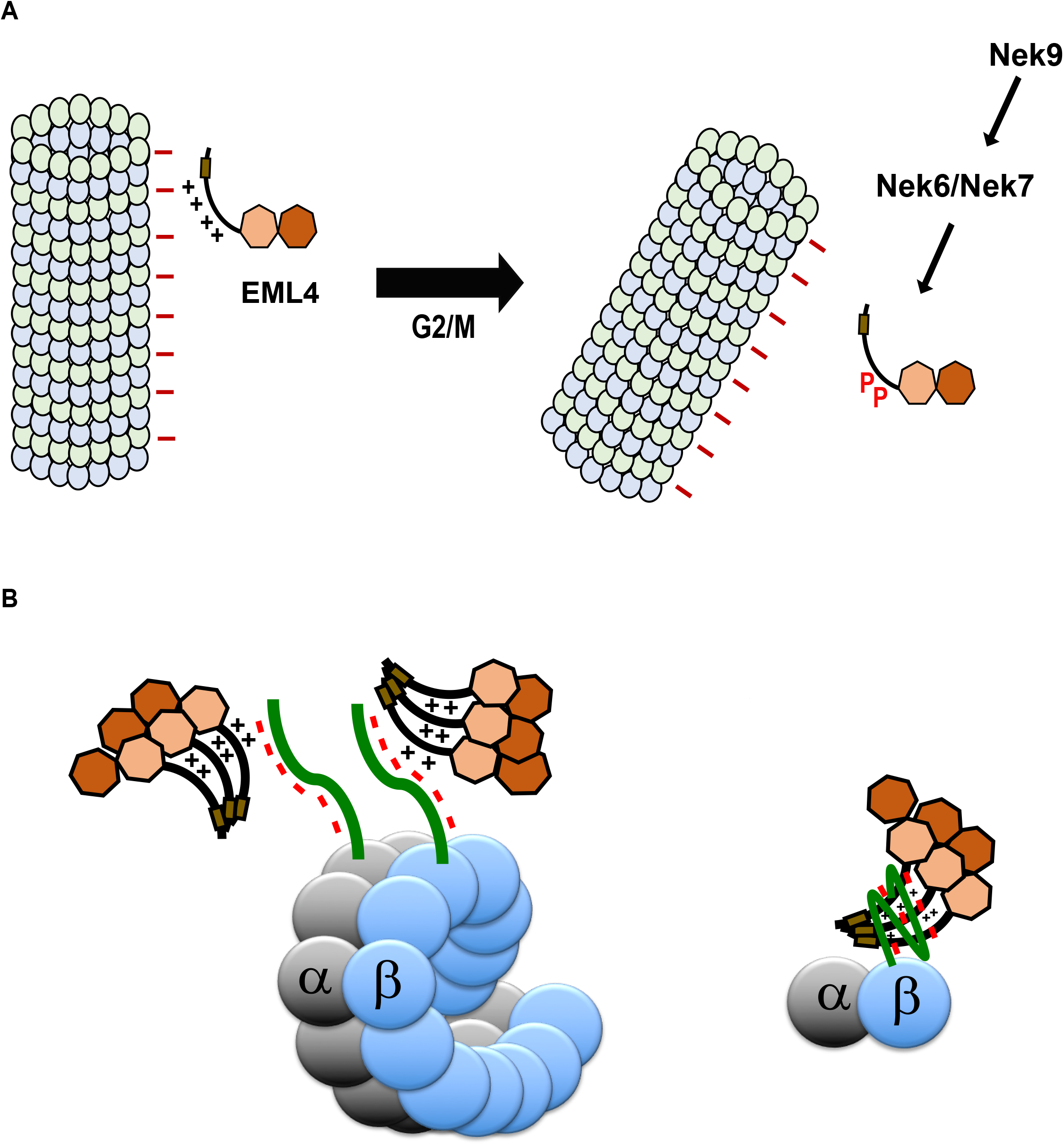
Phospho-dependent regulation of EML4 microtubule binding affinity. (**A**) Schematic model proposing how the Nek9-Nek6-Nek7 kinase module regulates affinity of EML4 for microtubules at the G2/M transition through phosphorylation of two sites within the EML4 NTD thereby reducing the electrostatic interaction of this basic region for the acidic microtubule surface. (**B**) Schematic cartoon showing potential mechanism of interaction of the EML4 NTD with tubulin C-terminal tails.

The mechanism by which EML4 stabilizes microtubules in cells, either directly or indirectly, also remains enigmatic. However, EML4 shares many features with the ch-TOG (XMAP215) family of MAPs in that both proteins have separable domains for binding the microtubule lattice and soluble tubulin. ch-TOG acts as a processive microtubule polymerase by binding to the microtubule with a basic region and then using its multiple TOG domains to add soluble tubulin to the growing microtubule plus-ends (Al-Bassam & Chang, 2011). However, although EML4 has a basic NTD that binds the microtubule polymer and a TAPE domain that binds soluble tubulin, there was no detectable concentration of EML4 at plus ends of microtubules where it could promote growth through acting as a microtubule polymerase. Interestingly, like tau, EML4 is abundant in the nervous system suggesting that it may have a major function in stabilizing the long microtubules present in neurons (Houtman et al., 2007). At the molecular level, proteins of the tau family bind to more than one tubulin dimer, thereby preventing catastrophe, slowing shrinkage and promoting rescue, while also being displaced from the microtubule lattice by phosphorylation (Bowne-Anderson et al., 2015, van der Vaart et al., 2009). The fact that EML4 exists as a trimer would similarly allow it to bind to more than one tubulin dimer, thereby increasing its overall avidity for the microtubule surface, and potentially directly stabilising the polymer. However, in contrast to tau (Kellogg, Hejab et al., 2018), EML4 does not interact precisely with the microtubule surface but, rather, loosely and flexibly associates with the tubulin C-terminal tails. Also in contrast to tau and its relatives, EML2 and the echinoderm EMAP have been reported to destabilise microtubules by increasing the catastrophe rate and decreasing the rescue rate, respectively (Eichenmuller et al., 2002, Hamill et al., 1998). In this regard, it is interesting that the non-neuronal isoform of EML2 used in this study, as well as EMAP, lack the trimerization domain raising the possibility that trimerization is necessary for EMLs to stabilise microtubules. Conversely, a longer isoform of EML2 that does have a trimerization domain is found in the brain and spinal cord where it could also stabilize microtubules (Lepley, Palange et al., 1999).

Another key finding of this study is the discovery of a new role for the mitotic Nek6 and Nek7 kinases. Depletion of Nek6 or Nek7, or the upstream kinase Nek9 that activates Nek6 and Nek7 upon mitotic entry, leads to increased association of EML4 with spindle microtubules. This provides compelling evidence that these enzymes play a central role in regulating the microtubule binding affinity of EML4. This hypothesis is supported by the observation that expression of constitutively active mutants of Nek6, Nek7 or Nek9 reduces binding of EML4 to microtubules in interphase. Although in theory this regulation could be indirect, we found that Nek6 and Nek7 were capable of phosphorylating EML4 in vitro, including at S144 and S146, while mutation of these two sites or depletion of Nek6, Nek7 or Nek9 led to loss of a gel-mobility shift of the EML4-NTD in mitosis. Together with the fact that EML4 is present in a complex with Nek6 and Nek7 in cells (Ewing et al., 2007), this strongly suggests that EML4 is a direct physiological substrate for these kinases. Depletion of Nek6 or Nek7 disrupts spindle formation and both the kinesin Eg5 and the chaperone Hsp72 have been reported as mitotic substrates of Nek6 (O’Regan et al., 2015, Rapley, Nicolas et al., 2008). Our work here indicates that EML4 should be added to that growing list of substrates for these kinases and that, like Cdk1, Plk1 and Aurora-A, the mitotic Nek kinases phosphorylate multiple substrates to drive robust spindle assembly. However, we do not rule out that EML4, and its related family members, may also be subject to phospho-dependent regulation by other mitotic kinases, with for example sea-urchin EMAP reported to be heavily phosphorylated by Cdk1(Brisch et al., 1996).

Removal of EML4 from microtubules therefore appears to be crucial for the decrease in stability that enables assembly of a highly dynamic mitotic spindle. Live cell imaging demonstrated that the ability of condensed chromosomes to congress to the metaphase plate was severely hampered in cells expressing a phosphonull mutant. This was accompanied by a delay in anaphase onset that was presumably mediated by persistent activity of the spindle assembly checkpoint. Overexpression of a wild-type EML4 protein also delayed chromosome congression and anaphase onset albeit to a much reduced extent compared to the phosphonull mutant. However, this would be consistent with the endogenous Nek6 and Nek7 kinases struggling to phosphorylate the full complement of excess EML4 protein. Interestingly, EML4 is nevertheless required for mitotic progression arguing that, despite the reduced microtubule binding affinity, it still plays an important role at this stage of the cell cycle. EML4 was found to bind the nuclear distribution C (NUDC) protein via its TAPE domain and target NUDC to the spindle promoting kinetochore capture by microtubules (Chen et al., 2015). Whether this requires the microtubule binding activity of EML4 is not known. Of course, while microtubule stability initially drops upon mitotic entry, specific populations of microtubules, notably K-fibres, become highly stabilised upon bi-orientation of chromosomes. Indeed, residual staining of spindle microtubules was detected and it is therefore possible that EML4 could contribute to K-fibre stability if a localized fraction of EML4 was dephosphorylated.

Many proteins specifically interact with microtubules during mitosis. These fulfil a variety of important functions that include the regulation of microtubule stability, the trafficking of checkpoint complexes, and the motility of the condensed chromosomes (Ferreira, Pereira et al., 2014). Collectively, these proteins enable the assembly of a dynamic and finely tuned mitotic spindle. In contrast, EML4 preferentially interacts with microtubules during interphase, and the regulated disruption of this interaction through phosphorylation is what is necessary for proper mitotic spindle assembly and function. This raises the question of how many other proteins might dissociate from microtubules during mitosis. Indeed, it seems likely that the electrostatic interactions that other microtubule-associated proteins depend on might also be disrupted through phosphorylation, as protein phosphorylation peaks during this phase of the cell cycle. This mode of regulation could therefore be relevant to other proteins, including potentially other members of the EML family, that interact with the negatively charged tubulin C-terminal tails.

## MATERIALS AND METHODS

### Plasmid construction, mutagenesis and recombinant protein expression

Generation of YFP-EML4-FL and YFP-EML1-NTD constructs was previously described(Richards et al., 2015), while YFP-EML4-NTD and YFP-EML4-TAPE were generated by PCR-based amplification of the NTD and TAPE domain fragments from the YFP-EML4-FL plasmid and subcloning into pLEICS-12 (PROTEX, University of Leicester). Generation of Flag-Nek6-Y108A, Flag-Nek7-Y97A, and Flag-Nek9-ΔRCC1 constructs were as described (O’Regan & Fry, 2009, Richards et al., 2009, Roig et al., 2002). Mutations were introduced into the YFP-EML4-FL and NTD constructs using the GeneTailor Site-Directed Mutagenesis Kit (Invitrogen), and all constructs confirmed by Sanger sequencing at the University of Leicester. The YFP-EML1-NTD and YFP-EML4-NTD proteins were expressed and purified as described (Richards et al., 2015), while Flag-Strep-EML4-FL protein used for phosphomapping was expressed in insect cells and purified as described(Richards et al., 2014).

### Cell culture, transfection and drug treatment

U2OS, HeLa and HEK 293 cells were grown in Dulbecco’s modified Eagle’s medium (DMEM, Invitrogen) supplemented with 10% heat-inactivated foetal bovine serum (FBS), 100 IU/ml penicillin and 100 µg/ml streptomycin at 37°C in a 5% CO_2_ atmosphere. HeLa Kyoto H2B-mCherry/EGFP-Lamin A cells were maintained in DMEM containing 10% FBS, 100 IU/ml penicillin, 100 µg/ml streptomycin, 500 µg/ml G418 and 0.5 mg/ml puromycin. Transient transfections were performed with Lipofectamine 2000 (Invitrogen) according to manufacturer’s instructions. Cells were synchronized in M-phase either by incubation for 16 h with 500 ng/ml nocodazole, or by incubation with 10 µM RO-3306 (Enzo Life Science) for 16 h followed by transfer into fresh media with 20 µM MG132 (EMD Millipore) for 2 h. M-phase arrested cells were collected by mitotic shake-off after 16 h treatment.

### Western blotting

Cells were lysed in ice-cold RIPA or NEB lysis buffer (50 mM Tris-HCl pH 8, 150 mM NaCl, 1% v/v Nonidet P-40, 0.1% w/v SDS, 0.5% w/v sodium deoxycholate, 5 mM NaF, 5 mM β-glycerophosphate, 30 µg/ml RNase, 30 µg/ml DNase I, 1x Protease Inhibitor Cocktail, 1 mM PMSF) and subjected to SDS-PAGE and analysis by Western blotting. Primary antibodies were rabbit Nek6 (1 µg/ml; (32), goat Nek7 (1:250; Aviva Systems), rabbit Nek9 (0.4 µg/ml; Atlas Antibodies), mouse α-tubulin (0.3 µg/ml; Sigma), mouse acetylated tubulin (1:2000; Sigma), rabbit GAPDH (1:500; Cell Signaling), rabbit EML4 (N-term, A301-908A; C-term, A301-909A; both 1:500, Bethyl Laboratories), rabbit green fluorescent protein (GFP; 0.5 µg/ml; Abcam), mouse pHistone H3 (1:1000, Abcam) and mouse cyclin B1 (0.5 µg/ml; Santa Cruz). Secondary antibodies were horseradish peroxidase (HRP)-labelled secondary antibodies (1:1000; Amersham).

### RNAi

Cells at 30–40% confluency were cultured in Opti-MEM Reduced Serum Medium with 10% heat-inactivated foetal bovine serum (FBS), and transfected with 50 nM ON-TARGETplus siRNA duplexes using Oligofectamine (Invitrogen) according to manufacturer’s instructions. siRNA duplexes were as previously described for Nek6 and Nek7 (O’Regan & Fry, 2009), Nek9 (AM51334-1113, Ambion) and EML4 (HSS120688 and HSS178451, Dharmacon). Cells were fixed or lysed for analysis after 72 h transfection.

### In vitro kinase assay and mass spectrometry

Kinase assays were carried out using 5-10 µl immunoprecipitates or 0.1 µg purified Nek6 or Nek7 kinase (Millipore). Proteins were incubated with 5 µg substrate and 1 µCi of [γ-32P]-ATP in 40 µl kinase buffer (50 mM Hepes.KOH pH 7.4, 5 mM MnCl_2_, 5 mM β-glycerophosphate, 5 mM NaF, 4 µM ATP, 1 mM DTT) at 30°C for 30 min. Reactions were stopped with 50 µl of protein sample buffer and analysed by SDS-PAGE and autoradiography. Phosphomapping was performed using an LTQ-Orbitrap-Velos-ETD (ThermoFisher Scientific) as previously described(O’Regan et al., 2015).

### Fluorescence microscopy

For immunofluorescence microscopy, cells were grown on acid-etched glass coverslips, fixed with ice-cold methanol or methanol: acetone (1:1) and processed as previously described (O’Regan & Fry, 2009). In brief, media were aspirated and cells fixed with ice-cold methanol at −20°C for 30 min. Cells were blocked in 1x PBS supplemented with 3% BSA and 0.2% Triton X-100 before incubation with antibodies in 1x PBS supplemented with 3% BSA. Primary antibodies used were mouse α-tubulin (0.3 µg/ml; Sigma-Aldrich), rabbit Flag (1:1000; Sigma-Aldrich), rabbit green fluorescent protein (GFP) (1 µg/ml; Abcam), rabbit EML4 (N-term, A301-908A; C-term, A301-909A; both 1:500, Bethyl Laboratories), mouse acetylated tubulin (1:2000; sigma), goat Flag (1:1000; Abcam), and rat α-tubulin conjugated 647 (1:200; Abcam). Secondary antibodies used were Alexa Fluor 488, 594 and 647 donkey anti-rabbit, donkey anti-mouse and donkey anti-goat IgGs (1 µg/ml; Invitrogen). DNA was stained with 0.8 µg/ml Hoechst 33258. Fixed and time-lapse imaging were performed on a Leica TCS SP5 confocal laser scanning microscope fitted on a Leica DMI 6000B inverted microscope using a hcx plan apo 63x oil objective (numerical aperture, 1.4). For fixed images, Z stacks comprising 30–50 0.3 µm sections were acquired using LAS-AF software (Leica), and deconvolution of 3D image stacks performed using Huygens Essential software (Scientific Volume Imaging). For time-lapse fluorescence imaging, cells were cultured in glass-bottomed culture dishes (MatTek Corporation) and maintained on the stage at 37°C in an atmosphere supplemented with 5% CO_2_ using an environmental chamber (Life Imaging Services). Z-stacks comprising of 0.5 µm sections were acquired every 10 min for a minimum of 16 h. Stacks were processed into maximum intensity projections using LAS-AF software (Leica) and movies prepared using ImageJ. To quantify colocalization, ImageJ software was used to draw five lines across the cytoplasmic regions of a cell on a single z-section of an image. A total of 10 cells from three independent experiments were used to calculate the mean Pearson’s correlation coefficient (R-value).

For structured illumination microscopy (SIM), cells were grown on acid-etched high performance coverslips, fixed in ice-cold methanol and stained for immunofluorescence microscopy with EML4 and α-tubulin antibodies as described above. However, following incubation with the secondary antibody, cells were subjected to a second, post-fixation step in which cells were incubated with 4% paraformaldehyde for 10 minutes, prior to coverslips being mounted onto microscope slides with Citifluor mounting medium (Electron Microscopy Sciences). Imaging was performed using a Zeiss PS1 super resolution microscope and images processed using Zen software (Zeiss).

For TIRF imaging, microtubules were prepared from a mixture of 2 µM biotin-labelled, 2 µM Hilyte647-labelled and 20 µM unlabelled tubulin, 1 mM GMP-CPP in MRB80 (80 mM K-PIPES, 4 mM MgCl_2_, 1 mM EGTA, pH 6.9). Microtubules were then treated with subtilisin (25 µg/ml) or left untreated and incubated for 20 min at 37°C. To terminate digestion 2 mM PMSF was added and the reaction mix centrifuged for 5 min at 150,000 xg. The supernatant was removed and microtubules re-suspended in MRB80 buffer. Microtubules were attached to the surface of a flow cell using PLL-PEG-biotin and streptavidin, and binding of 100 nM recombinant EML1 (1-174) was assessed using TIRF microscopy as described (Richards et al., 2015).

### In vitro microtubule binding and sedimentation assays

Microtubules were assembled from 40 µM tubulin in the presence of 5 mM GTP in MRB80 for 1 h at 37°C, diluted 1:3 in MRB80 + 2 µM Taxol. 3/5 of the sample was treated with 25 µg/ml subtilisin (Sigma P5380) and the remainder left untreated and incubated at 37°C for 10 min. To terminate digestion 5 mM PMSF and complete protease inhibitors (Roche) were added and the reaction mix loaded onto a 30% sucrose cushion and centrifuged in a TLA55 rotor at 100,000 xg for 45 min at 30°C. The microtubule pellets were re-suspended in BRB25 (25 mM PIPES pH 6.8, 1 mM MgCl_2_, 1 mM EGTA, 2 mM DTT) supplemented with complete protease inhibitors and 2 µM Taxol and incubated with EML4(1-207)-Avi on ice for 15 min before pelleting through a 30% Sucrose cushion in a TLA55 rotor at 100,000 xg for 45 min at 4°C. The pellet was taken up in equal volume to supernatant and samples analysed on 12% SDS-PAGE gels, stained with Instant Blue (Expedion), imaged on a G-Box (Syngene) and quantified using ImageJ.

For microtubule sedimentation assays, cells were lysed at room temperature in RIPA lysis buffer, including 5 mM NaF, 5 mM β-glycerophosphate. Samples were loaded on the top layer of 30% sucrose in tubulin stabilization buffer (TSB; 1 mM EGTA, 5 mM MgCl_2_, 80 mM PIPES at pH 7.0) before centrifugation at 100,000 g for 40 min at 21°C. Supernatants were collected and pellets washed with TSB for 10 min at 21°C. Pellets were diluted into a volume equal to that of the supernatants. Samples were subjected to SDS-PAGE and analysis by Western blot.

### Sample preparation for cryo-electron microscopy

Lyophilized tubulin purified from HeLa cells was purchased from Cytoskeleton Inc. (Denver, CO, USA) and reconstituted to 2.5 mg/ml in BRB80 (80 mM PIPES, 1.5 mM MgCl_2_, 1 mM EGTA, 1 mM DTT) containing 1 mM GMPCPP at 4°C. After 10 min incubation at 4°C the tubulin was transferred to a water bath and polymerised at 37°C for 45 min. In order to increase GMPCPP occupancy, microtubules were double-cycled by pelleting at 13,000 rpm on a desktop centrifuge at room temperature, removing the supernatant and re-suspending the microtubule pellet to ~2.5 mg/ml at 4°C in BRB80 + 1 mM GMPCPP. Microtubules were then repolymerised by incubation at 37°C for 45 mins. Stabilised microtubules were left at room temperature for 3 hours then diluted in room temperature BRB25 + 1 mM GMPCPP to 0.5 mg/ml before use (25 mM PIPES, 1.5 mM MgCl_2_, 1 mM EGTA, 1 mM DTT). 4 µl of microtubules were pre-incubated on glow-discharged holey C-flat™ carbon EM grids (Protochips, Morrisville, NC) at room temperature for 1 min, excess buffer manually blotted away, then 4 µl of 2.2 mg/ml EML4-NTD in BRB25 + 30 mM NaCl and 1 mM GMPCPP added for 45 sec. Excess buffer was again manually blotted away, followed by another 4 µl application of EML4-NTD. In an additional step for cryo-electron tomography only, excess buffer was again blotted away after 45 sec incubation and 4 µl of 10 nm nanogold fiducial– BSA solution (Sigma), concentrated 2-fold in BRB25 + 30 mM NaCl and 1 mM GMPCPP, added to the grid. Grids were then placed in a Vitrobot Mark IV (FEI Co., Hillsboro, OR) at room temperature and 80% humidity, incubated for a further 45 sec, then blotted and vitrified in liquid ethane.

### Single-particle cryo-electron microscopy data collection and processing

Low dose movies were collected automatically using EPU software on a K2 summit direct electron detector (Gatan) installed on a FEI Titan Krios (Astbury Biostructure Laboratory, University of Leeds) operating at 300kV with a quantum post-column energy-filter (Gatan), operated in zero-loss imaging mode with a 20-eV energy-selecting slit. A defocus range of 0.5-2.5µm and a calibrated final sampling of 1.37Å/pixel was used with the K2 operating in counting mode at 6e-/pixel/second. The total exposure was 48e-/Å^2^ over 30 frames (1.6e-/Å^2^/frame). Movie frames were aligned using Motioncorr2 (Zheng, Palovcak et al., 2017) with a patch size of 5 to generate full dose and dose-weighted sums. Full dose sums were used for CTF determination in gCTF (Zhang, 2016), then dose-weighted sums used in particle picking, processing and generation of the final reconstructions. Particle processing was performed with helical methods in Relion v2.1.0 (He & Scheres, 2017), using a custom pipeline and scripts to account for MT pseudo-helical symmetry. Input references were 7 Å low-pass filtered density maps of undecorated 13 protofilament GMPCPP microtubules made from protofilaments of the undecorated 14 protofilament GMPCPP microtubule atomic model (PDB: 6DPU (Zhang, LaFrance et al., 2018)). Final displayed reconstructions were sharpened to local resolutions as determined in Relion, unless stated. The atomic models of 6 dimers of undecorated GMPCPP MTs (PDB: 6DPU) with incorporated taxol (taken from PDB: 5SYF (Kellogg, Hejab et al., 2017)) were fitted into symmetrized density and used as starting models for iterative rounds of model building in Coot (Emsley & Cowtan, 2004) and Phenix (Afonine, Poon et al., 2018). Extra density contributed by EML4-NTD had low resolution, presumably due to the flexible nature of the MT-EML4-NTD interaction; therefore EML4-NTD was not modelled into density.

### Cryo-electron tomography data collection and processing

Single-axis cryo-electron tomography of EML4-NTD decorated microtubules was performed using a Tecnai G2 Polara at 300 kV with a Quantum post-column energy filter (Gatan) operated in zero-loss imaging mode with a 20-eV energy-selecting slit. Data at 5–6 µm defocus were collected on a K2 Summit direct electron detector operating in counting mode at 9e-/pixel/second (measured without sample obstructing the beam) with a final sampling of 5.39Å per pixel. Tilt series of total dose 114e-/Å^2^ from −60 to +60° tilt were collected in 3° increments using the Hagen dose-symmetric tilt scheme (Hagen, Wan et al., 2017). For each tilt, movies of 9 seconds exposure at four subframes per second were aligned using MotionCor2 (Zheng et al., 2017). Dose weighting of tilt series was performed using custom scripts calling functions in SumMovie (Grant & Grigorieff, 2015). Fiducial-based alignment of tilt series was performed in the Etomo graphical user interface to IMOD (v.4.9.0). CTF determination on each aligned tilt series without dose-weighting was performed with CTFFIND4 (Rohou & Grigorieff, 2015) and three-dimensional CTF correction and tomogram reconstruction was performed by weighted back-projection of dose-compensated and aligned tilt series with novaCTF (Turonova, Schur et al., 2017). Final tomograms displayed were 4xbinned using IMOD and a B-factor of 35000 applied to amplify low-frequency information.

### Statistical analysis

All quantitative data represent means and standard deviation of at least three independent experiments. Statistical analyses were performed using a one-tailed unpaired Student’s t test assuming unequal variance; *, *p*<0.05; **, *p*<0.01; ***, *p*<0.001. n.s., non-significant.

## ACKNOWLEDGMENTS

This work was funded by grants from Worldwide Cancer Research (16-0119) and The Wellcome Trust (204801/Z/16/Z) to A.M.F., the Medical Research Council, U.K. (MR/R000352/1) to C.A.M, Cancer Research UK (C24461/A23302) to R.B., and The Wellcome Trust (200870/Z/16/Z) and Lister Institute of Preventive Medicine to A.S. We acknowledge support from the University of Leicester Core Biotechnology Services (CBS) for DNA sequencing, construct preparation and confocal imaging, and Robert Markus at the School of Life Sciences Imaging Facility at the University of Nottingham for super-resolution microscopy. Single-particle cryo-EM data was collected with Rebecca Thompson at the Astbury Biostructure Laboratory, University of Leeds. The FEI Titan Krios microscopes were funded by the University of Leeds (UoL ABSL award) and Wellcome Trust (108466/Z/15/Z). Cryo-EM structures have been deposited as EMD-0331, PDB ID 6I2I.

## AUTHOR CONTRIBUTIONS

R.A., J.M.M., J.A., L.O., A.S., R.B., C.A.M. and A.M.F. designed research; R.A., J.M.M., J.A., L.O. and D.R. performed research; R.A., J.M.M., L.O. and M.W.R. contributed new reagents; R.A., J.M.M., J.A., L.O., K.R.S., D.R., A.S., R.B., and A.M.F. analysed data; R.A., J.M.M., J.A., L.O., A.S., R.B., C.A.M. and A.M.F. wrote the paper.

## SUPPLEMENTARY INFORMATION

**Supplementary Figure S1.**
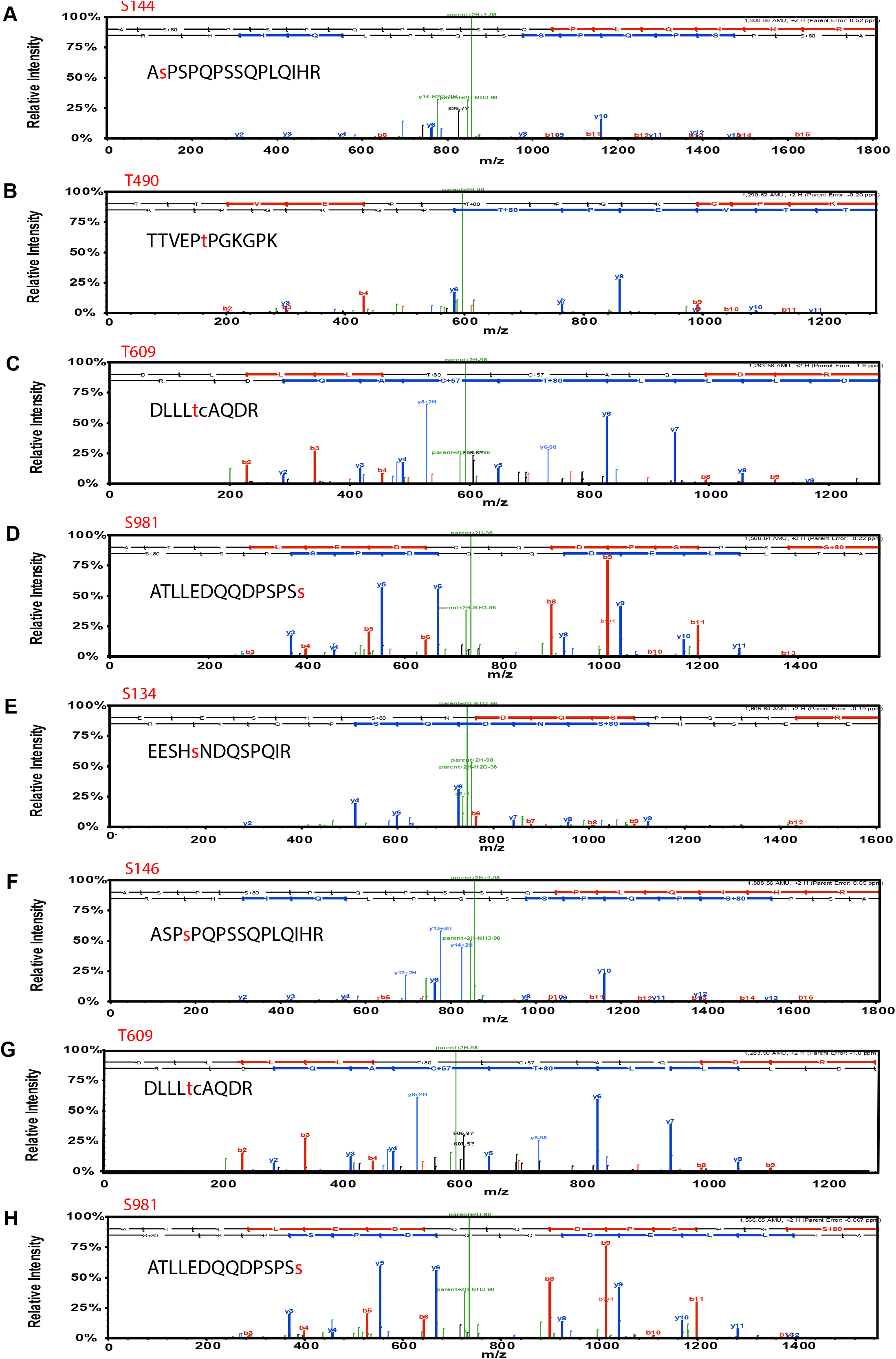
Mass spectrometry profiles of EML4 phosphorylation sites. Purified EML4 was phosphorylated with NEK6 and NEK7, in vitro. Mass spectrometry profiles were categorised as NEK6 (A-D) and NEK7 (E-H). Phosphorylated amino acids are indicated in red in the identified peptide.

**Supplementary Figure S2.**
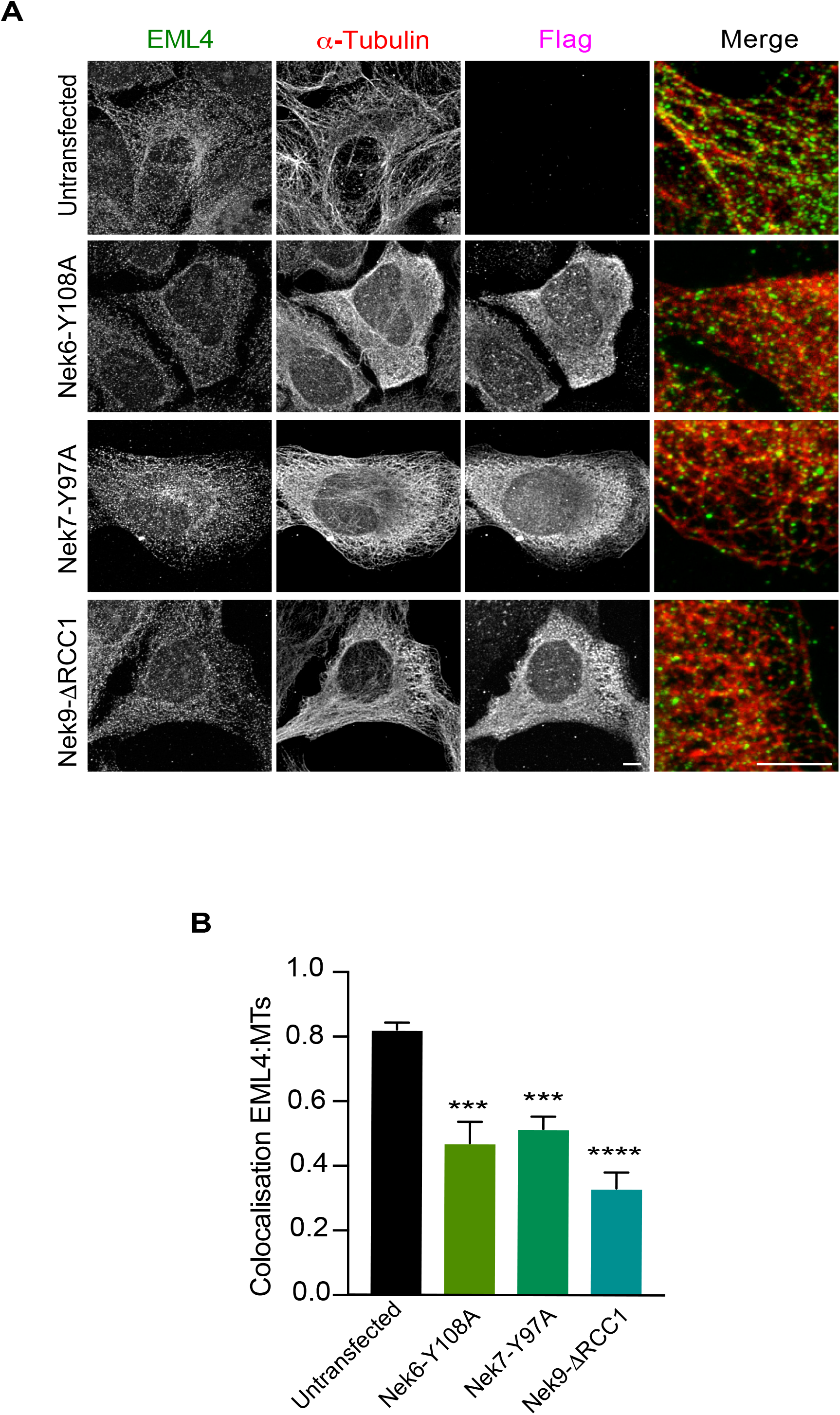
Activated Nek6, Nek7 and Nek9 reduce association of EML4 with interphase microtubules. (**A**) U2OS cells were mock-transfected or transfected with Flag-Nek6-Y108A, Flag-Nek7-Y97A, or Flag-Nek9-ΔRCC1 for 24 h before being stained with EML4, α-tubulin and Flag antibodies. (**B**) The mean Pearson’s correlation coefficient for co-localization between EML4 and microtubules was calculated from 5 lines per cell in 10 cells ±SD.

**Supplementary Figure S3.**
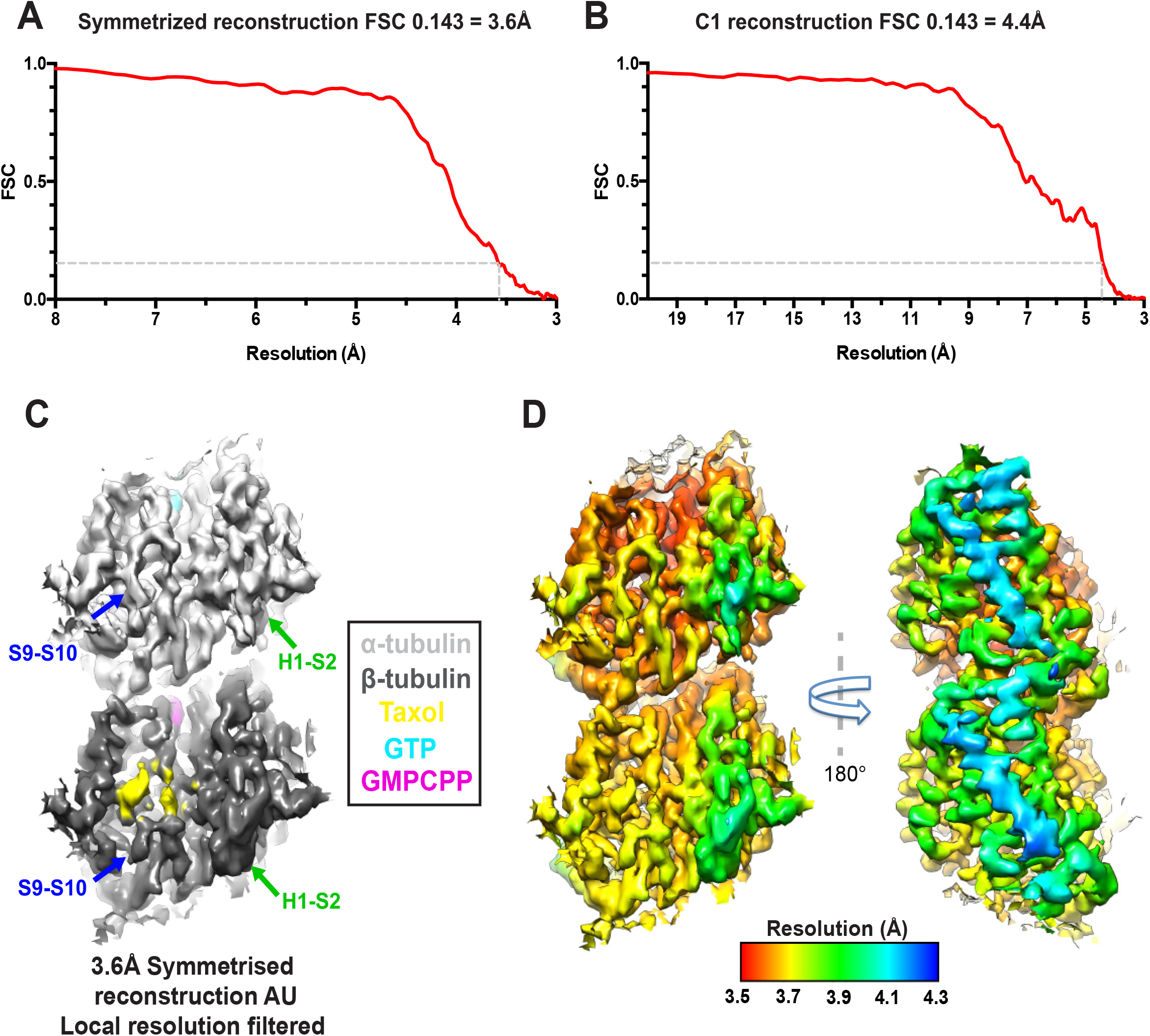
Cryo-electron microscopy diagnostics and resolution estimation. (**A**) Gold-standard FSC curve for the symmetrized 13pf EML4-MT reconstruction. (**B**) Gold-standard FSC curve for the C1 13pf EML4-MT reconstruction. The central 15% of the microtubule along the helical axis was isolated from the independently refined half-maps (using a soft-mask) for FSC calculations. (**C**) An asymmetric unit in the symmetrized reconstruction viewed from the MT lumen, showing α- and β-tubulin in light and dark grey respectively. Densities for GTP, GMPCPP and paclitaxel (included in the tubulin purification protocol) are indicated in cyan, magenta and yellow, respectively. Density for the S9-S10 and H1-S2 loops exhibit expected differences in α- and β-tubulin, indicating successful seam determination during the image alignment. Because of the reference used for alignment, the central, best region of the reconstruction spans across an inter-dimer longitudinal contact encompassing α- and β-tubulin from separate dimers. (**D**) Local resolution calculated in Relion for an asymmetric unit of the symmetrized reconstruction, showing lumenal (left) and outer (right) faces. A B-factor of −90 was applied to sharpen the reconstructions shown in panels C and D, up to local resolution cut-offs as displayed in panel D.

**Supplementary Figure S4.**
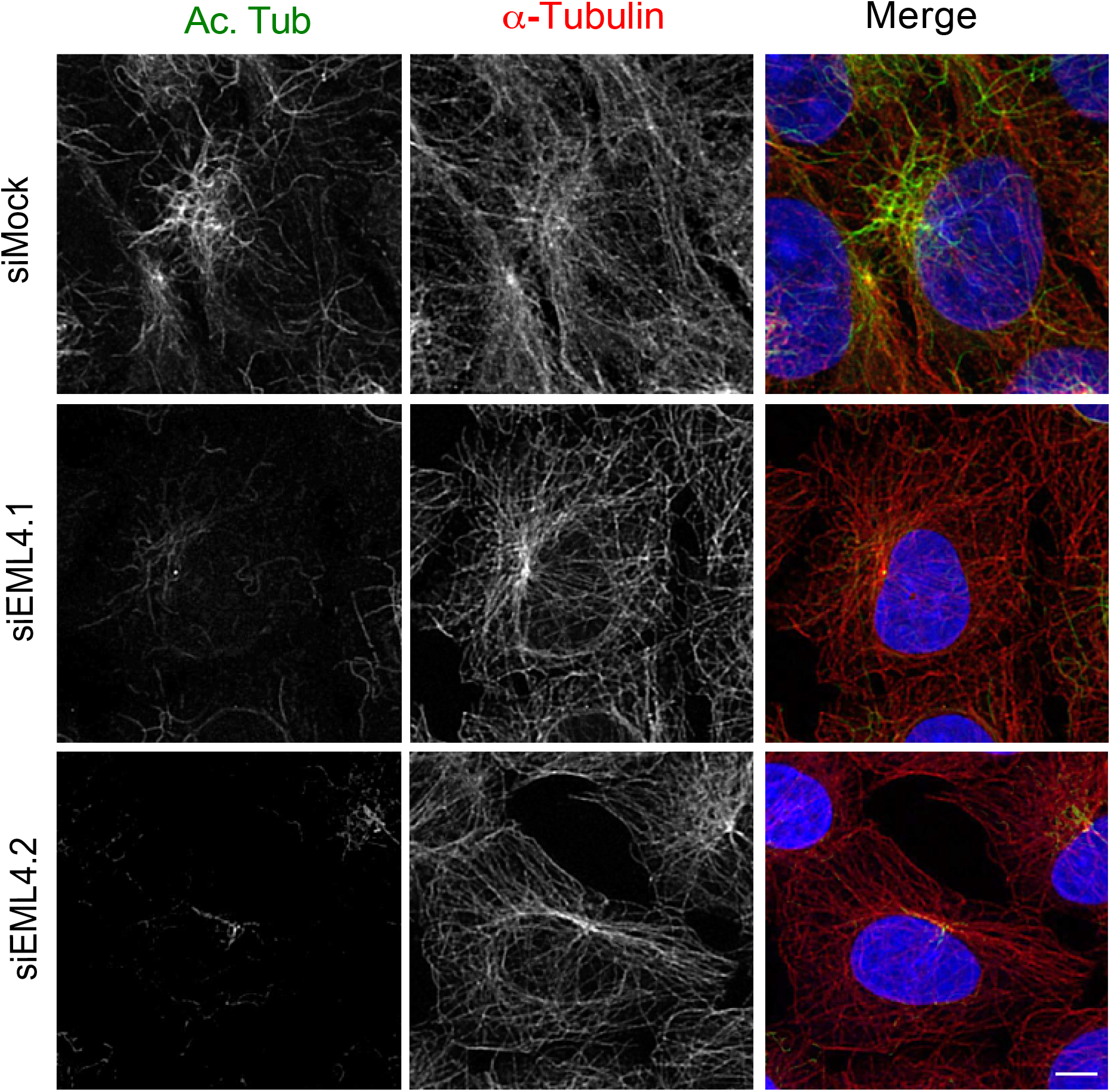
EML4 depletion leads to reduced microtubule acetylation. U2OS cells were either mock or EML4-depleted for 72 h before staining with acetylated tubulin (green) and α-tubulin (red) antibodies. Merge images include DNA stained with Hoechst 33258 (blue). Scale bar, 5 µm.

